# SSRIs modulate asymmetric learning from reward and punishment

**DOI:** 10.1101/2020.05.21.108266

**Authors:** Jochen Michely, Eran Eldar, Alon Erdman, Ingrid M. Martin, Raymond J. Dolan

**Affiliations:** Max Planck UCL Centre for Computational Psychiatry and Ageing Research, University College London, London, United Kingdom; Wellcome Centre for Human Neuroimaging, University College London, London, United Kingdom; Department of Psychiatry and Psychotherapy, Charité – Universitätsmedizin Berlin, Berlin, Germany; Psychology and Cognitive Sciences Departments, Hebrew University of Jerusalem, Jerusalem, Israel; Institute of Cognitive Neuroscience, University College London, London, United Kingdom

## Abstract

Human instrumental learning is driven by a history of outcome success and failure. We demonstrate that week-long treatment with a serotonergic antidepressant modulates a valence-dependent asymmetry in learning from reinforcement. In particular, we show that prolonged boosting of central serotonin reduces reward learning, and enhances punishment learning. This treatment induced learning asymmetry can result in lowered positive and enhanced negative expectations. A consequential effect is more rewarding, and less disappointing, experiences and this may, in part, explain the slow temporal evolution of serotonin’s well-established antidepressant effects.

## Introduction

To make good choices, agents need to learn from past success and failure (Skinner, 1938). Human learning is characterised by remarkable flexibility, including adjustment to environmental volatility (Behrens *et al*., 2007; McGuire *et al*., 2014), and an asymmetric impact of positive and negative reinforcement (Daw *et al*., 2002; Gershman, 2015). The latter is assumed to play a role in the emergence of mood disorders, often characterised in terms of aberrant processing of reward and punishment (Murphy *et al*., 2003; Pulcu & Browning, 2017). However, it remains unclear whether learning asymmetries represent biases contributing to suboptimal reward prediction, or adaptive computations supporting risk and threat avoidance necessary for survival (Niv *et al*., 2012; Caze & van der Meer, 2013).

Learning from reinforcement is attributed to two key components: a prediction error (PE), quantifying a difference between expectation and outcome, and a learning rate, determining the impact of PEs on future reward predictions (Sutton & Barto, 1998). There is evidence that PEs also impact emotional states (Rutledge *et al*., 2014; Eldar & Niv, 2015; Eldar *et al*., 2018), where mood depends not only on how well things are going in general but whether things are better or worse than expected. Based on the latter findings it follows that learning asymmetries can, in principle, exert a modulatory effect on mood. For instance, enhanced learning from reward generates a high expectation of future reward, limiting subsequent opportunities for the experience of momentary happiness arising out of positive surprise. Similarly, slow learning from punishment can potentially lead to repeated disappointment due to negative surprise, leading to an emergence of low mood (Eldar *et al*., 2016b; Kaye & Ross, 2017).

In this study, we tested a hypothesis of an asymmetric effect of serotonergic antidepressants on reinforcement learning that can account for a positive impact on mood. We used computational modelling to assess learning from reward and punishment in a sample of healthy volunteers exposed to week-long treatment with selective serotonin reuptake inhibitors (SSRIs), a first-line antidepressant intervention (Hieronymus *et al*., 2018). Specifically, we conjectured SSRIs would selectively impact on learning asymmetries, reducing learning from reward, and enhancing learning from punishment, in a manner that results in an accumulation of more positive and less negative surprise, respectively.

## Results

66 healthy volunteers were administered either a daily oral dose of the SSRI citalopram (20mg) or placebo across seven consecutive days. Subjects performed two experimental sessions, once after administration of a single dose on day 1 and once, after exposure to repeated daily administration, on day 7 (Fig. 1B). On each session, subjects performed a modified version of a gambling card game (Fig. 1A; (Eldar *et al*., 2016a)), where the goal was to maximize monetary wins and minimize monetary losses.

**Figure 1.**
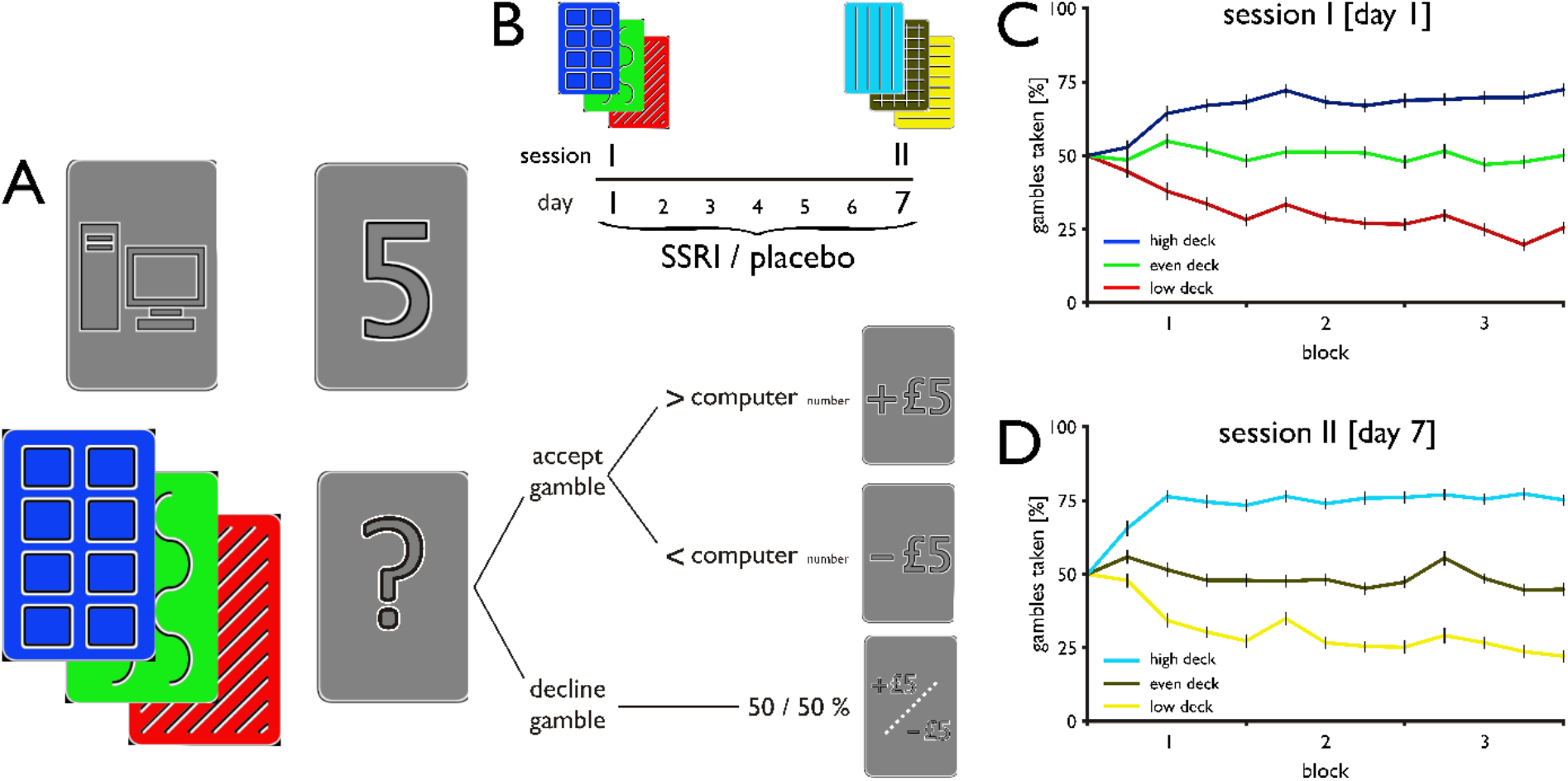
Experimental task, pharmacological procedure, and learning performance. **(A)** *Experimental design*. On each trial, participants were presented with one of three possible decks and a number between 1 and 9 drawn by the computer. If participants decided to gamble, they won £5 if the number they drew was higher than the computer drawn number, and lost £5 if the number was lower. Participants were only informed whether they won or lost the gamble, not which number they drew. Participants had to learn by trial and error how likely gambles were to succeed with each of the three decks. One deck contained a uniform distribution of numbers between 1 and 9 (even deck), one deck contained more 1’s (low deck), making gambles 30% less likely to succeed, and one deck contained more 9’s (high deck), making gambles 30% more likely to succeed. Opting to decline the gamble resulted in a 50% probability of win/loss, regardless of which number was drawn by the computer. **(B)** *Pharmacological procedure*. Subjects were randomly allocated to take a daily dose of 20mg citalopram or placebo for seven days and participated in two sessions: session I took place on day 1 after single administration, session II took place on day 7, at a time when citalopram reaches steady-state plasma levels. Subjects played an identical game on both sessions, but with two independent sets of three decks, where colour order and colour-associated win probability randomly varied across participants. **(C) & (D)** *Gambles taken with each deck as a function of time*. Percentages were computed separately for each set of 15 contiguous trials (4 sets/60 trials per block), for session I **(C)**, and session II **(D)**, respectively.

In brief, on each trial participants were presented with a number between 1 and 9 as drawn by a computer. Subjects could gamble that the number they were about to draw would be higher than the computer drawn number. Critically, participants played with one of three possible decks on each trial, which differed in how likely gambles were to succeed. One deck contained a uniform distribution of numbers between 1 and 9 (even deck), one deck contained more 1’s (low deck, gambles 30% less likely to be successful), and one deck contained more 9’s (high deck, gambles 30% more likely to be successful). Subjects were informed that an unsuccessful gamble would result in a loss (−£5), and a successful gamble would result in a win (+£5). Subjects learnt through trial and error about each of the decks’ success likelihood. Alternatively, subjects could decline the gamble to opt for a fixed 50% known probability of winning or losing, respectively. On the second session, the game was identical, and the only difference being that subjects played with three novel decks, indicated by different colours, where colour order and colour-associated win probability was randomly varied across participants (Fig. 1B). Subjects had to learn about these decks anew as they were unrelated to the ones from the first session.

On both sessions, subjects’ willingness to gamble differed depending on each deck’s win likelihood as a function of time (Fig. 1C/D). A trial-by-trial logistic regression confirmed subjects, on both sessions, gambled more against lower computer numbers (P<0.001), and gambled more with better decks (P<0.001), consistent with successful trial-and-error learning (Fig. 2A/C). Additionally, over the course of a session, participants gambled more with accumulating reward (P<0.001), and avoided gambling with accumulating punishment (P<0.001). On session I, effects were similar across drug groups for successful (rewarded), and unsuccessful (punished) gambles (P=0.41 & P=0.36; Fig. 2B). At session II, however, we found evidence for an asymmetric impact of reward and punishment, as a function of treatment (drug x valence, P=0.02; Fig. 2D), and this was attributable to an enhanced impact of losses (P=0.01), but not wins (P=0.31), in SSRI as compared to placebo subjects. This differential pattern suggests prolonged SSRI treatment enhanced gambling avoidance as a function of cumulative punishment.

**Figure 2.**
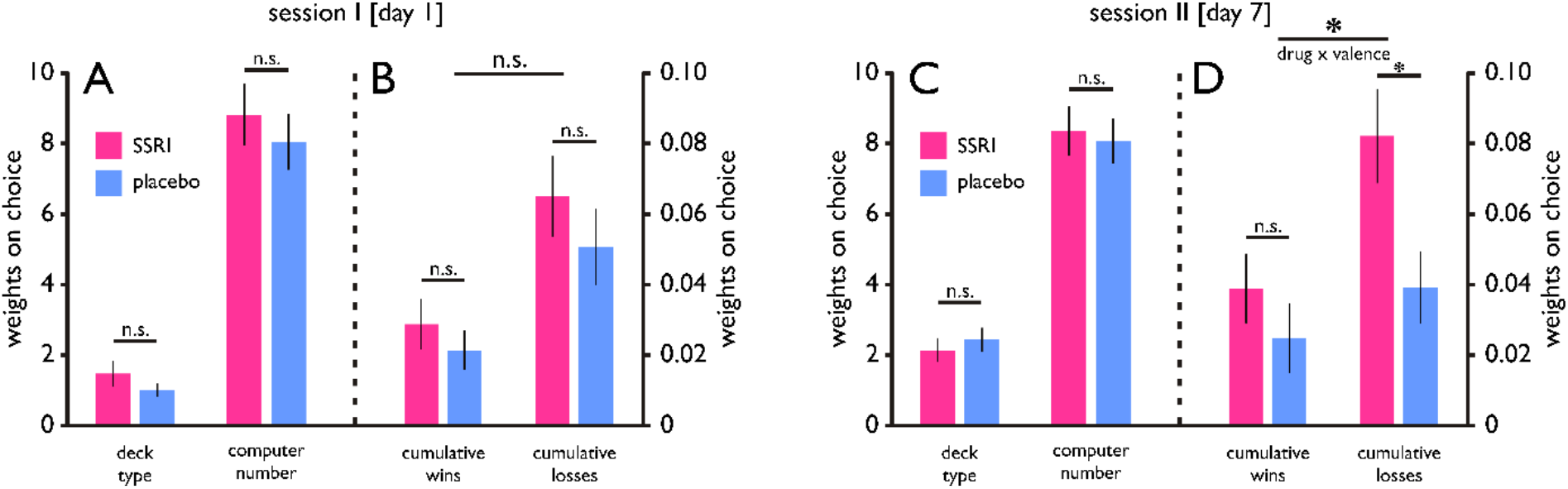
Results of trial-by-trial logistic regression model. **(A) & (C)** Fitting a logistic regression model to subjects’ decisions showed that participants gambled more with better decks (deck type: −1 for low, 0 for even, 1 for high) and against lower computer numbers (scaled to range between −1 (for number 9) and 1 (for number 1), with no drug differences on session I **(A)**, and session II **(C)**, respectively. **(B) & (D)** Additionally, subjects, over the course of a session, gambled more with increasing rewards (cumulative wins for each deck, reflecting the sum of previous positive gamble outcomes, computed as +1 multiplied by the computer’s number against which it was received), and gambled less with increasing punishment (cumulative losses for each deck, reflecting the sum of previous negative gamble outcomes, computed as −1 multiplied by (10 – the computer’s number against which it was received). On session I, impact of preceding wins and losses was unaffected by treatment **(B)**. On session II, however, SSRIs enhanced the impact of losses but not wins **(D)**, indicating an asymmetric drug effect on reward and punishment.

In principle, participants can acquire information about decks both from success and failure. To account for the precise mechanisms guiding learning, we used computational modelling. Replicating results from an earlier study using an identical cognitive task (Eldar *et al*., 2016a), model comparison showed behaviour was best explained by a model that accounted for an asymmetry in learning from the two outcome types. Specifically, the best-fitting model included two different learning rates, one for wins (η^+^), and one for losses (η^-^), where these determine the degree to which an outcome type impacts on subsequent expectations (model 6: ‘*adjusted & asymmetric Q-learning*’; Supplementary file 1A). These expectations, in combination with the numbers drawn by the computer, shape whether gambles are likely to be taken or declined.

On session I, learning from reward and punishment was similar across treatment groups (η^+^: P=0.70, η^-^: P=0.67 Fig. 3A). However, by session II, there emerged a significant serotonergic impact on learning asymmetry (drug x valence, P=0.007; Fig. 3B), such that, in SSRI as compared to placebo subjects, learning from reward was reduced (P=0.009), while learning from punishment was enhanced (P=0.04).

**Figure 3.**
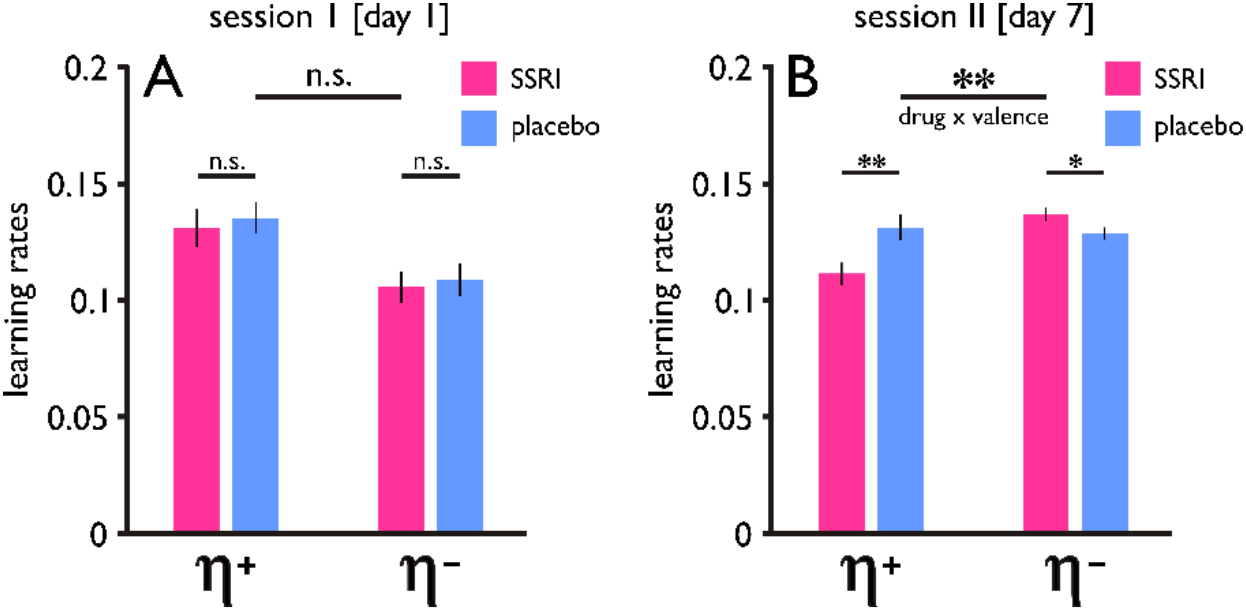
Learning asymmetry and its serotonergic modulation. **(A)** On session I, computational modelling showed learning from reward (η^+^) and learning from punishment (η^-^) was unaffected by drug treatment. **(B)** On session II, SSRI treatment had a significant effect on learning asymmetries, such that it reduced learning from reward (η^+^), and enhanced learning from punishment (η^-^).

Additionally, an asymmetric effect of cumulative reward and punishment on gambling, as derived from the logistic regression, correlated significantly with the asymmetry in learning derived from our computational model (session I: r=0.76, P<0.001, session II: r=0.63, P<0.001; Fig. 3–Fig. supplement 2). This indicates that an altered gambling preference, related to altered learning from different outcome types, was a consequence of temporally extended serotonergic intervention.

We found a significant asymmetric effect on learning rates across data from both sessions (drug x valence: P=0.04), but no significant three-way interaction (drug x valence x session: P=0.34, controlling for an overall gambling bias, Fig. 3–Fig. supplement 1). There were no between-group differences for the remaining model parameters (Fig. 3–Fig. supplement 1). Lastly, generating simulated data based upon model parameters derived from the fitting procedure showed that the model captured core features of the real data (Fig. 2–Fig. supplement 1), individual parameter estimates could be accurately recovered (Supplementary file 1B/C), and our model comparison procedure could identify the model that generated the data (Supplementary file 1D).

## Discussion

Studies probing a serotonergic modulation of human reinforcement learning, using serotonergic depletion (Seymour *et al*., 2012), or single dose SSRI administration (Skandali *et al*., 2018), have proven inconclusive (Boureau & Dayan, 2011; Faulkner & Deakin, 2014). Here, we provide evidence that a temporally extended treatment with SSRIs exerts opposing effects on reward and punishment learning.

We suggest the asymmetric effect we highlight can go some ways to explain a gradual evolution of an incremental impact of SSRIs on mood. Recent studies have shown that positive and negative surprise strongly impacts on self-reported affective states (Rutledge *et al*., 2014; Eldar & Niv, 2015; Eldar *et al*., 2018). Given a key role of surprise, our results explain how serotonergic intervention can, in principle, influence the affective experience of reinforcement. Specifically, SSRIs give rise to more positive surprises by slowing down reward learning, and minimize subsequent disappointments by enhancing punishment learning. This computational mechanism may thus lead to a gradual emergence of better mood by virtue of an overall greater sampling of positive surprise.

Although a three-way interaction was not significant, evidence suggests that the effects we highlight require more extended treatment in order to evolve. Non-human animal data similarly show that a modulation of serotonin levels impacts learning processes at different timescales, with distinct effects of acute and prolonged intervention (Bari *et al*., 2010). The fact that changes in learning emerge only after an extended intervention may reflect two processes, or a synergism of both. First, citalopram reaches steady-state plasma levels after seven days (Gutierrez & Abramowitz, 2000), and boosting of serotonin levels by single dose administration is unlikely to be sufficient to induce substantial modulation of learning processes. Second, synaptic plasticity that may underlie this modulation, such as changes in BDNF levels or autoreceptor expression, require days or weeks to emerge (Krishnan & Nestler, 2010). The temporal trajectory of the effects we describe also mimics the time period for an early onset of subtle symptom changes following SSRI treatment in those with clinical depression (Taylor *et al*., 2006). Our study was restricted to a sample of non-depressed healthy individuals. Longitudinal assessment of clinical cohorts is an important next step in exploring the relationship between a temporal evolution of learning asymmetries and the emergence of clinically significant antidepressant effects over time.

In summary, we show that week-long SSRI treatment reduces reward and enhances punishment learning. This learning asymmetry can, in theory, result in lowered positive and enhanced negative expectations, and consequentially, to more rewarding, and less disappointing experience. We suggest this modulation of computations that guide reinforcement learning may contribute to a known serotonergic impact on mood.

## Methods

### Subjects

66 healthy volunteers (mean age: 24.7±3.9; range 20-38 years; 40 females; Supplementary file 1E) participated in this double-blind, placebo-controlled study. All subjects underwent an electrocardiogram to exclude QT interval prolongation and a thorough medical screening interview to exclude any neurological or psychiatric disorder, any other medical condition, or medication intake. The experimental protocol was approved by the University College London (UCL) local research ethics committee, with informed consent obtained from all participants.

### Pharmacological procedure

Participants were randomly allocated to receive a daily oral dose of the SSRI citalopram (20mg) or placebo, over a period of seven consecutive days. All subjects performed two laboratory testing sessions. Session I was on day 1 of treatment, 3h after single dose administration, as citalopram reaches its highest plasma levels after this interval (Noble & Benfield, 1997). On the following days, subjects were asked to take their daily medication dose at a similar time of day, either at home or at the study location. Session II was on day 7 of treatment, a time when citalopram is known to reach steady-state plasma levels (Gutierrez & Abramowitz, 2000), with the tablet being taken 3h before test.

### Affective state questionnaires

To examine putative effects of the drug on subjective affective states over the course of the study, participants completed the Beck’s Depression Inventory (BDI-II, (Beck *et al*., 1996)), Snaith-Hamilton Pleasure Scale (SHAPS, (Snaith *et al*., 1995)), State-Trait Anxiety Inventory (STAI, (Spielberger *et al*., 1983)), and the Positive and Negative Affective Scale (PANAS, (Watson *et al*., 1988)) on two different occasions: (i) pre-drug, day 1; (ii) peak drug, day 7.

### Experimental task

To examine differences in learning from success and failure, we used a modified version of a gambling card game (Eldar *et al*., 2016a), in which subjects’ goal was to maximize monetary wins and minimize monetary losses.

The game consisted of 180 trials, divided into three 60-trial blocks. On each trial (Fig. 1A), subjects were shown with which one of three possible decks (each designated by distinct colour and pattern) they will be playing. After a short interval (2 to 5 s, uniformly distributed), the computer drew a number between 1 and 9, and participants had up to 2.5 s to choose whether they wanted to gamble that the number, which they are about to draw, will be higher than the computer drawn number. If participants chose to gamble, they won £5 if the number that they drew was higher than the computer’s number, and they lost £5 if it was lower (as well as in half of the trials in which the numbers were equal). If subjects opted to decline the gamble, they won/lost with a fixed 50% known probability. Not making any choice always resulted in a loss. Feedback was provided 700 ms following each choice and consisted of a ‘+£5’, ‘-£5’, or ‘+£5 / −£5’ visual symbol. The drawn number was not shown. Subjects were told that each of the three decks contained a different proportion of high and low numbers, and they could learn by trial and error about each of the decks’ likelihood of success.

Unbeknownst to participants, one deck contained a uniform distribution of numbers between 1 and 9 (‘even deck’), one deck contained more 1’s than other numbers (‘low deck’), making gambles 30% less likely to succeed, and one deck contained more 9’s than other numbers (‘high deck’), making gambles 30% more likely to succeed. In the first 15 trials, the computer drew the numbers 4, 5, and 6 three times each, and the other numbers once each. To make sure that all participants take a gamble in approximately 50% of trials, the numbers that the computer drew three times were increased by one (e.g., [4, 5, 6] to [5, 6, 7]), in each subsequent set of 15 trials, if subjects took two thirds or more of the gambles against these numbers in the previous 15 trials, or decreased by one if participants took a third or less of the gambles. Participants’ decks were pseudorandomly ordered while ensuring that the three decks were matched against similar computers’ numbers and that no deck appeared in successive trials more than the other decks.

On both sessions, the game was identical, with the only difference being subjects played with distinct sets of three decks, indicated by different colours, where colour order and colour-associated win probability randomly varied across participants (Fig. 1B). Subjects were informed that the decks from session II were entirely unrelated to the ones from session I, and they had to learn about the novel decks anew.

To familiarize participants with the basic structure of the task, subjects, on both sessions, performed a 60-trial training block with an ‘even’ deck, where visual feedback indicated the number that participants drew.

### Logistic regression analysis

We fitted a trial-by-trial logistic regression model to subjects’ decisions, comprising four different terms: (i) deck type (−1 for low, 0 for even and 1 for high), (ii) computer number (scaled to range between −1 (for number 9) and 1 (for number 1)), (iii) cumulative wins (reflecting, for each deck, the sum of previous positive outcomes, computed as +1 multiplied by the computer’s number against which it was received), and (iv) cumulative losses (reflecting, for each deck, the sum of previous negative outcomes, computed as −1 multiplied by (10 – the computer’s number against which it was received)).

### Computational modelling

#### Model space

To account for the precise mechanisms that guided learning from reward and punishment, we compared a set of computational learning models in terms of how well each model explained subjects’ choices. In all models, the probability of taking a gamble was modelled by applying a logistic function to a term that represented available evidence.

Model 1 (‘*gambling bias*’) and model 2 (‘*gambling bias & computer number*’) are oblivious to previous experience with the decks, and do not assume any learning to occur.

Here, model 1 computes the evidence as:

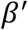
where *β*′ is a gambling bias parameter, determining a subject’s general propensity to gamble, thus allowing the model to favour either gambling or declining to begin with.

Model 2 computes the evidence as:

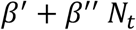
where N is the computer drawn number at trial t, scaled to range between −1 (for number 9) and 1 (for number 1), equivalent to the logistic regression, and *β*″ is an inverse temperature parameter, determining the strength, with which the computer’s number is determining a decision to gamble.

Model 3 (‘*Q-learning*’) learns the expected outcome of gambles with each deck as follows:

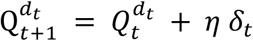
where

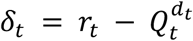

is an outcome prediction error, reflecting the difference between the actual (*r_t_*) and the expected 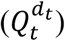 outcome of a gamble. *r_t_* = 1 represents a win, and *r_t_* = −1 represents a loss, and *η* is a learning rate parameter that weights the influence of prediction errors on subsequent expectations. Model 3 then computes the evidence as:

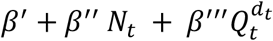
where *β*′′′ is a free parameter that determines the strength, with which choices are directed towards higher Q-value options.

In contrast to the previous model, model 4 (‘*adjusted Q-learning*’) computes prediction errors with respect to expectations that additionally factor in the computer’s number:

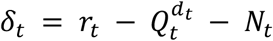
which means the model learns more from more surprising outcomes, i.e., from win outcomes of gambles against higher numbers, and from loss outcomes of gambles against lower numbers.

Based on prior work (Gershman, 2015; Eldar *et al*., 2016a), we assumed subjects would learn at a different rate from successful, i.e., reward, and unsuccessful gambles, i.e., punishment. In contrast to the gambling bias parameter (*β*′) that was included in all models, allowing them to favour either gambling or declining to begin with, an asymmetric learning bias can make such a tendency evolve with learning over time.

To this end, model 5 (‘*asymmetric Q-learning*’) and model 6 (‘*adjusted & asymmetric Q-learning*’) incorporate two distinct learning rate parameters (*η^+^* & *η*^-^), that allow learning at a different rate from different outcome types, i.e., from wins:

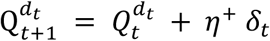
and from losses:

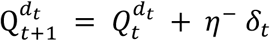

#### Model fitting

To fit the parameters of the different models to subjects’ decisions, we used an iterative hierarchical expectation-maximization procedure across the entire sample, separately for each session (Bishop, 2006; Eldar *et al*., 2018). We sampled 10^5^ random settings of the parameters from predefined prior distributions. Then, we computed the likelihood of observing subjects’ choices given each setting and used the computed likelihoods as importance weights to re-fit the parameters of the prior distributions. These steps were repeated iteratively until model evidence ceased to increase. To derive the best-fitting parameters for each individual subject, we computed a weighted mean of the final batch of parameter settings, in which each setting was weighted by the likelihood it assigned to the individual subject’s decisions. Note that the hierarchical fitting procedure, including all priors, were applied to the entire sample without distinguishing between SSRI and placebo subjects. This ensured that the parameter estimates, at the level of individual subjects, were mutually independent given the shared prior, rendering it appropriate to assess between-group differences. Learning rate parameters (*η*, *η*^+^ & *η*^-^) were modelled with Beta distributions (initialized with shape parameters *a* = 1 and *b* = 1). The gambling bias parameter (*β*′) was modelled with a normal distribution (initialized with *μ* = 0 and *σ* = 1), and inverse temperature parameters (*β*″ & *β*′′′) were modelled with Gamma distributions (initialized with *κ* = 1, *θ* = 1).

#### Model comparison

We compared between models in terms of how well each model accounted for subjects’ choices by means of the integrated Bayesian Information Criterion (iBIC (Huys *et al*., 2012; Eldar *et al*., 2018)). Here, we estimated the evidence in favour of each model (λ) as the mean likelihood of the model given 10^5^ random parameter settings drawn from the fitted group-level priors. We then computed the iBIC by penalizing the model evidence to account for model complexity as follows: iBIC = − 2 ln λ + κ ln n, where κ is the number of fitted parameters, and n is the total number of subject choices used to compute the likelihood. Lower iBIC values indicate a more parsimonious model fit.

## Acknowledgements

J.M. was supported by a fellowship from the German Research Foundation (MI 2158/1-1). R.J.D. holds a Wellcome Trust Investigator award (098362/Z/12/Z). The Max Planck UCL Centre for Computational Psychiatry and Ageing Research is a joint initiative supported by the Max Planck Society and University College London. The Wellcome Centre for Human Neuroimaging is supported by core funding from the Wellcome Trust (091593/Z/10/Z).

## Author contributions

J.M. and E.E. designed the experiments. J.M. and I.M.M. performed the experiments. J.M., E.E. and A.E. analysed and interpreted the data. J.M., E.E. and R.J.D. wrote the paper.

## Competing interest statement

The authors declare no competing interests.

## Data and code availability

Data and code for reproducing the associated analyses are available on GitHub: https://github.com/jmichely/ssri_asymmetric_learning.

---

**Figure 2 – figure supplement 1.**
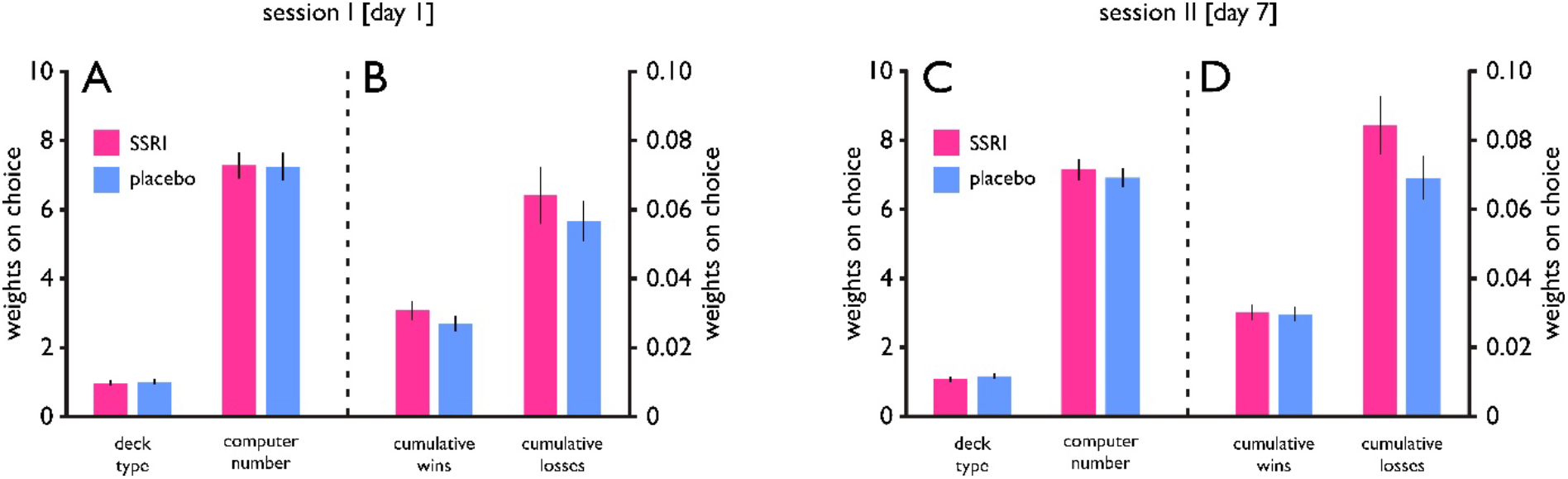
Results of trial-by-trial logistic regression model for simulated data. Generating simulated data based upon the model parameter estimates from the best-fitting model showed that the model captured core features of the real data. Note that for this analyses, we simulated 100 data sets and averaged the results. Fitting a logistic regression model to subjects’ simulated decisions revealed highly similar effects of **(A) & (C)** deck type and computer number, as well as **(B) & (D)** cumulative wins and losses on both sessions, respectively (cf. Fig. 2 of the main manuscript for the results of the real data). Error bars indicate SEM.

**Figure 3 – figure supplement 1.**
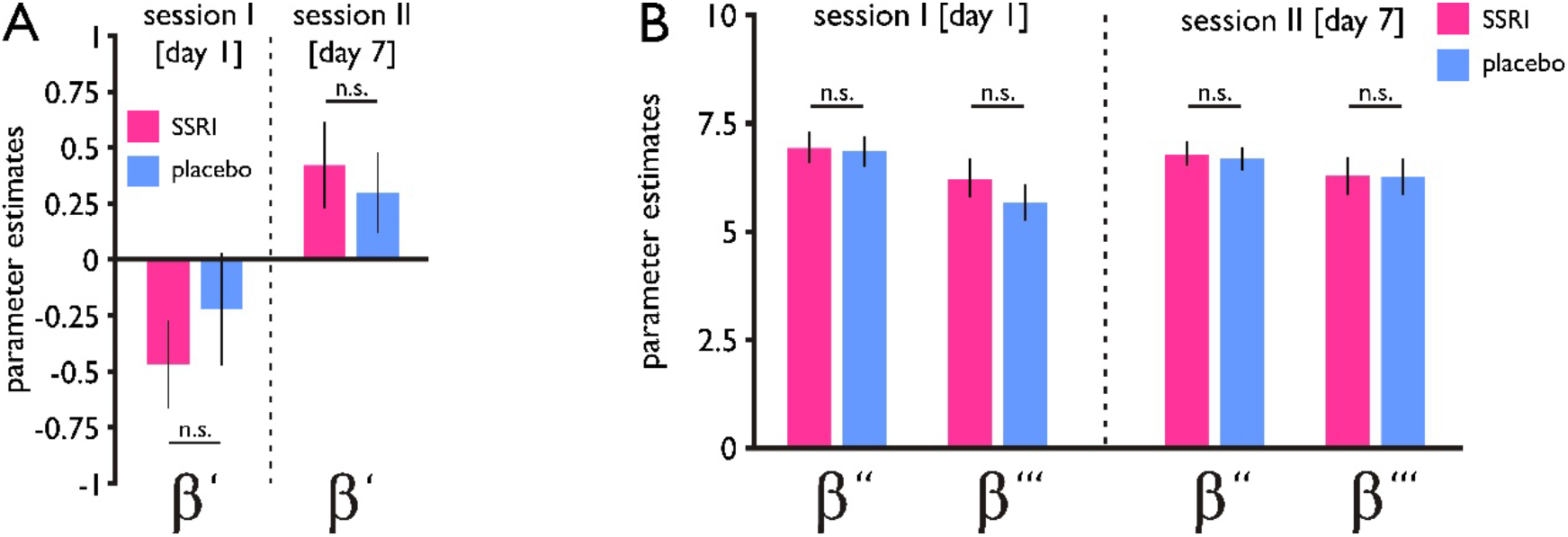
No drug effects on model parameters. In contrast to the effect of SSRI treatment on learning rates, there were no between-group differences for the remaining model parameters, such as **(A)** the gambling bias parameter (*β*′; session 1: P=0.87, session 2: P=0.78), and **(B)** the decision temperature parameter determining the impact of the computer number (*β*″; session1: P=0.40, session 2: P=0.97) and the impact of learned Q-values (*β*′′′ session1: P=0.44, session 2: P=0.64). n.s. = not significant. Error bars indicate SEM. Note that a gambling bias (*β*′) was overall negatively associated with a learning asymmetry (r=-0.18, P=0.03), in line with the notion that subjects with a positive gambling bias (more likely to gamble to begin with) have more room to adjust their behaviour through learning from negative events (negative learning asymmetry, η^-^ > η^+^).

**Figure 3 – figure supplement 2.**
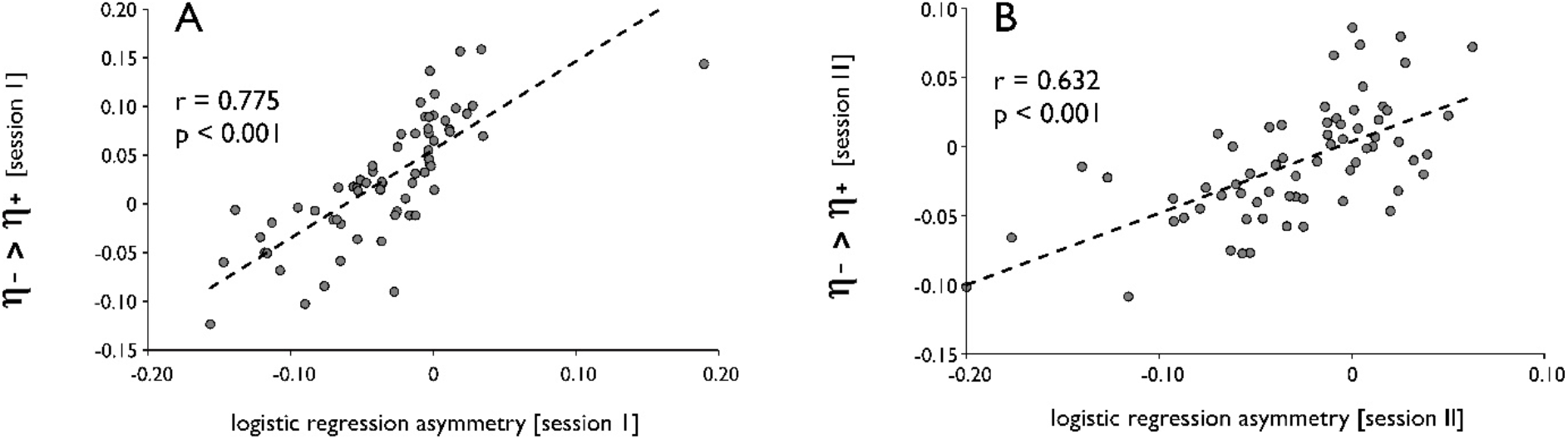
Asymmetric effects of reward and punishment. **(A)** An asymmetric effect of cumulative reward and punishment on gambling, as derived from the logistic regression, significantly correlated with an asymmetry in learning, as derived from our computational model, both on session I **(A)**, and on session II **(B)**.

## Supplementary File 1

**Supplementary file 1A.**
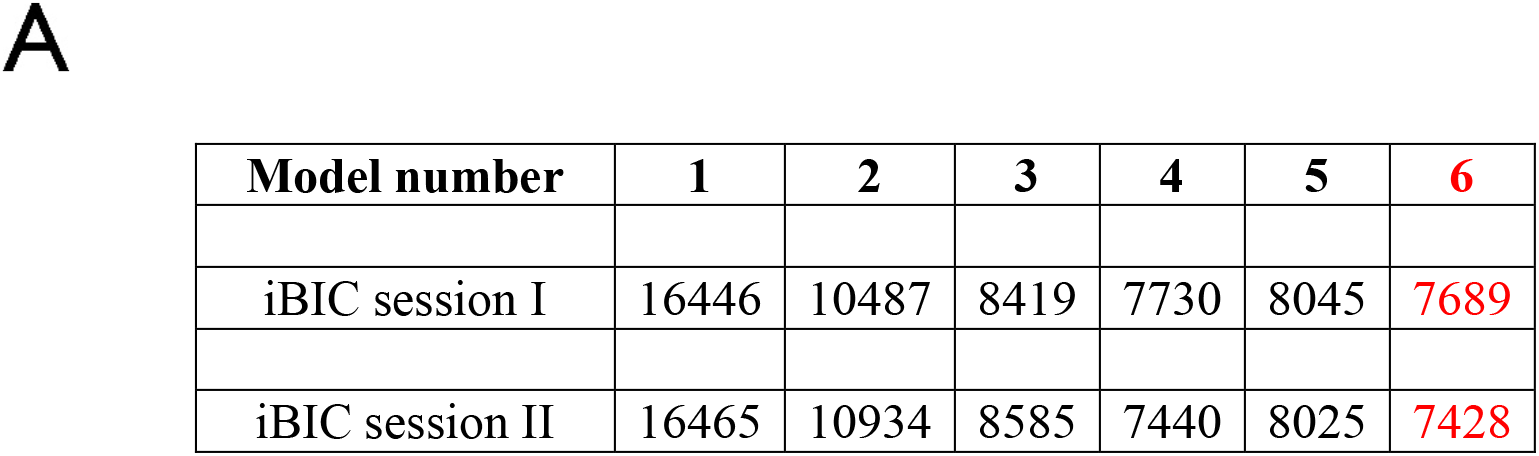
Model comparison. We compared 10 different models in terms of how well they explained subjects’ choices on each session. For each model, iBIC scores (integrated Bayesian Information Criterion) are shown. A lower iBIC score indicates better fit with subjects’ choices. The best-fitting model 6 is indicated in red. Model 1 (‘*gambling bias*’); Model 2 (‘*gambling bias & computer number*’); Model 3 (‘*Q-learning*’); Model 4 (‘*adjusted Q-learning*’); Model 5 (‘*asymmetric Q-learning*’); Model 6 (*‘adjusted & asymmetric Q-learning*’). Cf. Methods for details.S

**Supplementary file 1B/C.**
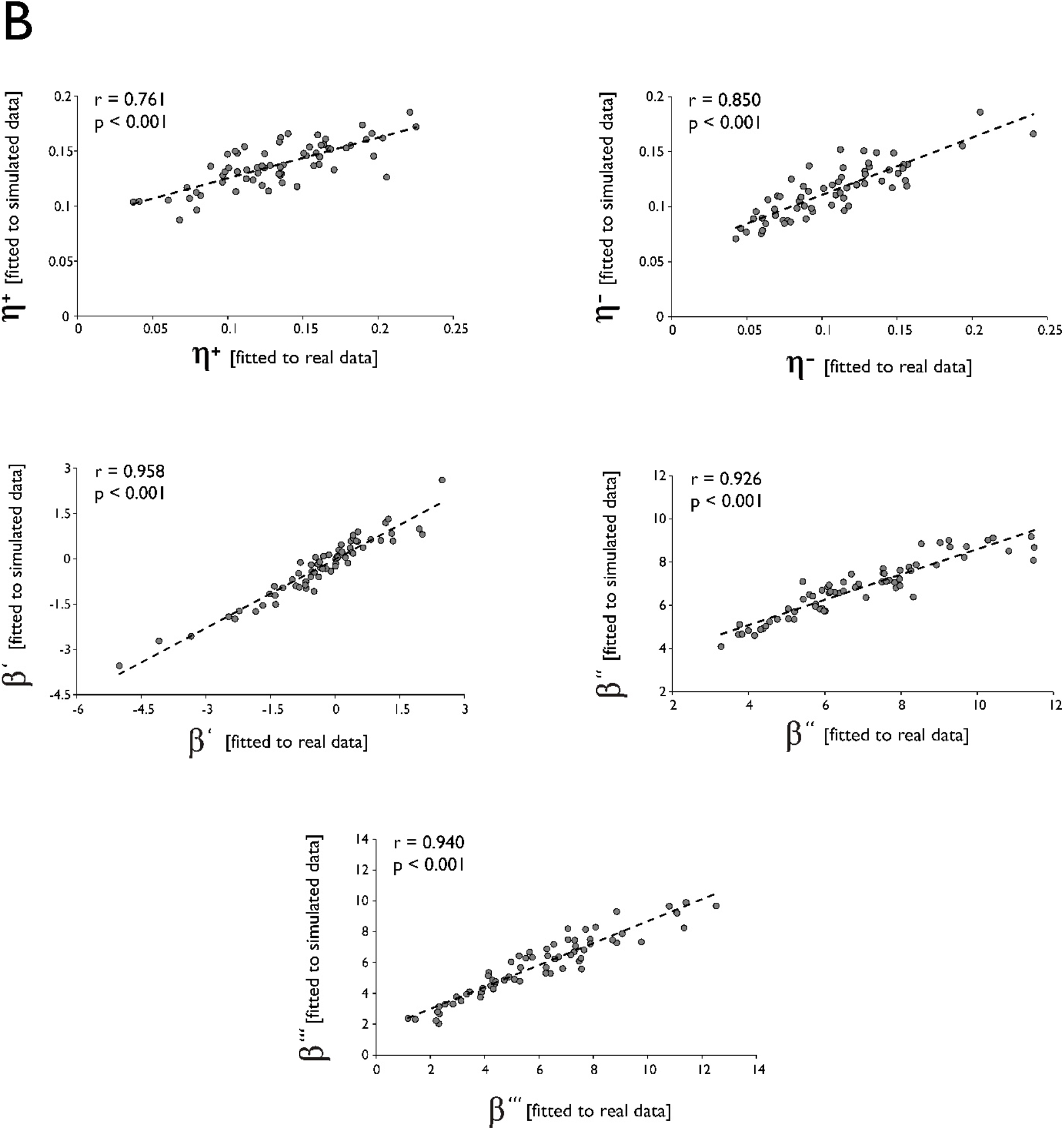
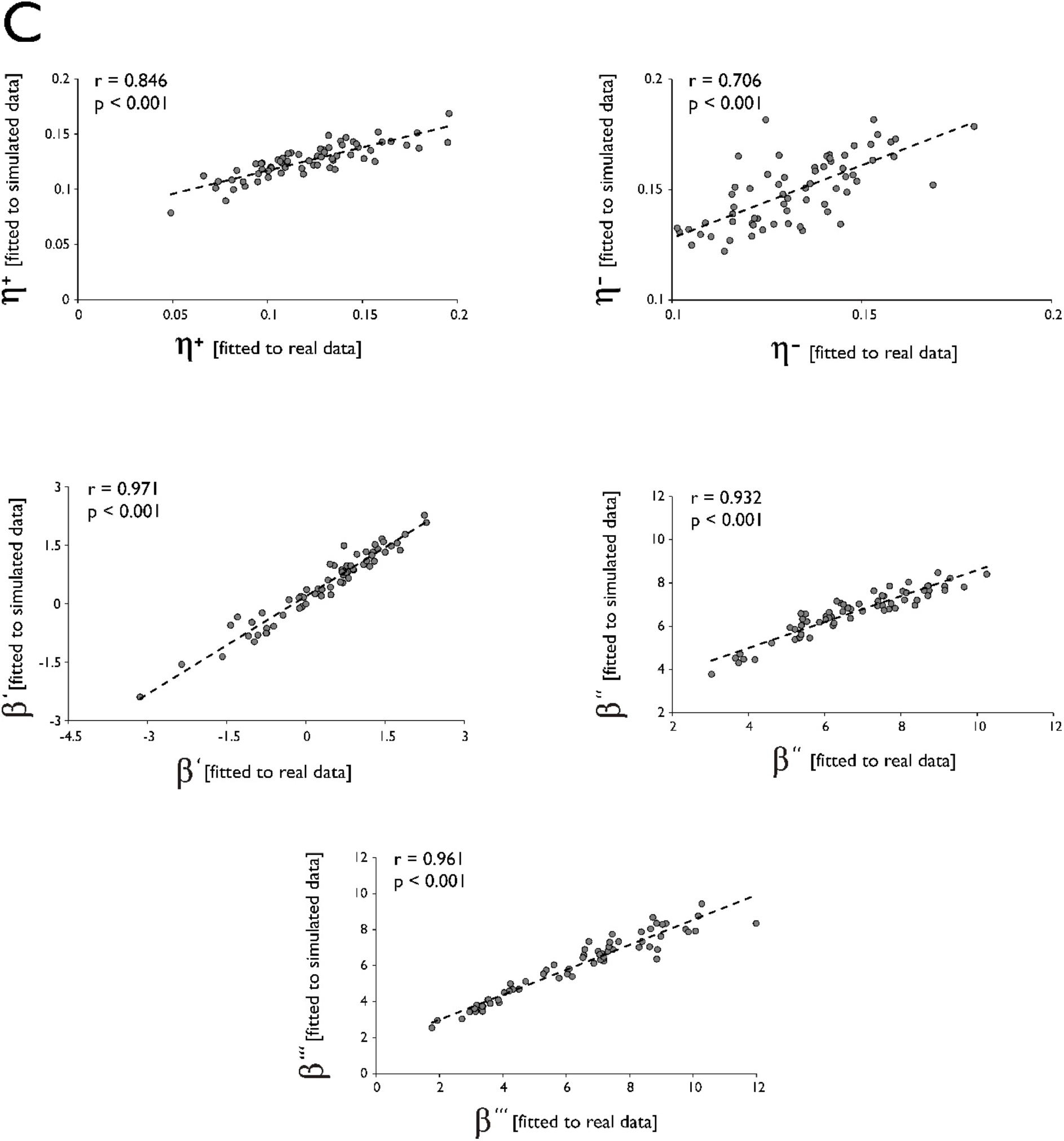
Recovery of model parameter estimates. Upon fitting the best-fitting model 6 to the simulated data, we found that model parameter estimates could be accurately recovered. This is indicated by a strong positive correlation between parameter estimates derived from fitting to real data (x-axis) and derived from fitting to simulated data (y-axis), both for **(A)** Session I, and **(B)** Session II. *β*′ = gambling bias parameter; *β*″ = decision temperature parameter determining the impact of the computer number; *β*′′′= decision temperature parameter determining the impact of learned Q-values; η^+^ = learning rate for reward; η^-^ = learning rate for punishment.

**Supplementary file 1D.**
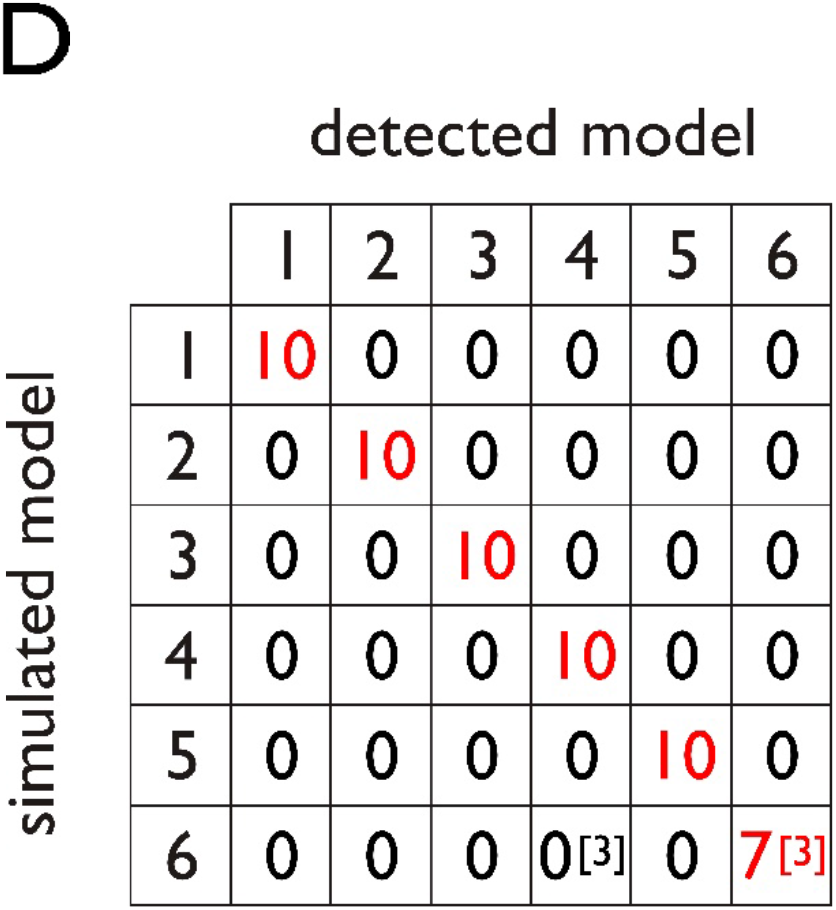
Validation of the model comparison procedure. We simulated 10 data sets across both sessions, using each model with its parameters fitted to subjects’ real choices, and we applied the model comparison procedure to each data set. Each cell shows how many datasets generated by the model indicated on the vertical axis were detected as reflecting the model used for simulation indicated on the horizontal axis. This analysis showed that the model comparison could accurately detect the model used for data simulation as the best-fitting model when using models 1-5 (10 out of 10 times, indicated in red), confirming specificity of the model comparison procedure, i.e., model 6 is not recognized when it is not the true underlying model. Moreover, this procedure confirmed sensitivity of the model comparison model procedure, i.e., model 6 is recognized when it is the true underlying model (10 out of 10 times). This was the case 7/10 times as the sole winner, i.e., iBIC difference > 6 to second best model, and 3/10 times as shared winner, i.e., within 6 of the lowest iBIC.

**Supplementary file 1E.**
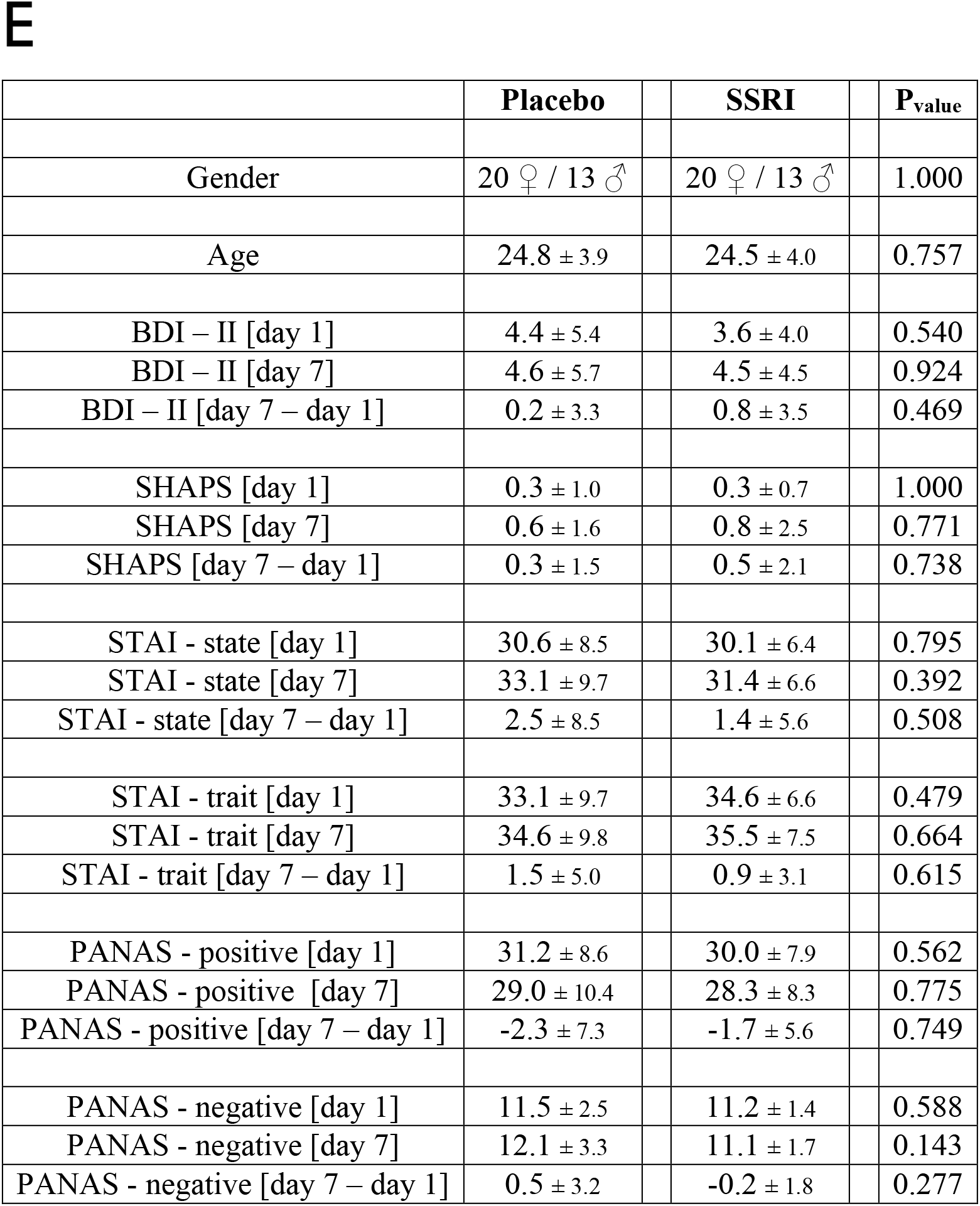
Affective state questionnaire data. Drug groups were matched for age and gender, and there was no baseline difference in any of the affective state questionnaires (assessed on day 1, pre-drug). Moreover, there was no drug effect on any of the affective state measures. BDI – II = Beck’s Depression Inventory II (Beck et al., 1996), SHAPS = Snaith-Hamilton Pleasure Scale (Snaith et al., 1995), STAI = State-Trait Anxiety Inventory (Spielberger, 1983), PANAS = Positive and Negative Affective Scale (Watson et al., 1988).

